# Matrix Stiffening Induces Mechanical Memory and Nuclear Fragility in Cardiomyocytes via Microtubule-Lamin Coupling

**DOI:** 10.1101/2025.07.17.665449

**Authors:** Nesrine Bouhrira, Sebastian Pizarro, Victor Senf, Kenneth B. Margulies

## Abstract

**Background:** Mechanical memory (MM) describes the persistent phenotypic remodeling following exposure to a transient extrinsic biomechanical cue. Short-term biomechanical stress is a feature of several etiologies of cardiomyopathy, including dysfunction of the viable myocardium following a large myocardial infarction. However, the nuclear mechanisms linking stiffness to persistent cellular remodeling remain poorly understood.

**Methods:** We cultured human iPSC-cardiomyocytes on a magnetorheological elastomer (MRE) with tunable stiffness (9–56 kPa) to mimic physiological and pathological myocardium. This allowed us to assess how transient increases in stiffness influence cellular responses such as nuclear structure, and DNA damage in hiPSC-cardiomyocytes. Using a combination of Immunofluorescence imaging, Western Blot and pharmacological interventions, we examined the role of microtubule detyrosination and the LINC (Linker of Nucleoskeleton and Cytoskeleton) complex in transducing mechanical signals from the cytoskeleton to the nucleus with a focus on MM induction.

**Results:** Short-term (6 h) stiff priming induced reversible phenotypic changes upon resoftening. However, 48 h of stiff priming triggered persistent MM, characterized by nuclear rupture, increased lamin A/C expression, DNA damage, and cytoplasmic leakage of DNA repair factors like KU80. Disruption of either a-tubulin detyrosination or the LINC complex prevented MM and nuclear damage, indicating that these elements are essential for nuclear mechanotransduction. In contrast, depletion of lamin A/C or DNA repair components accelerated stiffness-induced phenotypes and promoted MM onset within 6 hours. Finally, inhibition of a-tubulin detyrosination using ADV-TTL reversed both MM and DNA damage.

**Conclusions:** Acceleration of MM induction by lamin knockdown suggests that hereditary laminopathies may be associated with increased cardiomyocyte vulnerability to transient mechanical stress inducing DNA damage and senescence. Conversely, the protective effects of limiting stiffness-induced α-tubulin detyrosination, or nuclear mechano-transduction suggest potential cardioprotective strategies in the setting of laminopathies and/or sustained increases in extracellular matrix stiffness.

## Introduction

The nucleus is a dynamic mechanosensing organelle. In cardiomyocytes, extracellular mechanical cues are transmitted to the nucleus via the cytoskeleton, which physically connects to the nuclear envelope^1,2^. Forces acting upon the nuclear envelope, in turn, trigger changes in chromatin architecture and epigenetic regulators of gene expression^1,3–6^, which mediate mechano-responsive changes in cell size, shape, and function^7–9^. Temporal factors further modulate cardiomyocyte responses to mechanical cues^1^ [PMID: 37686151]. While short-term extracellular stiffening induces readily reversible phenotypic adaptations, more sustained exposure to relatively subtle increases in extracellular stiffening induces persistent changes in cellular structure and chromatin architecture. This phenomenon is referred to as "mechanical memory"(MM)^10^. Recent studies by our group demonstrated that immature cardiomyocytes derived from human induced pluripotent stem cells (hiPSC-CMs) exhibit persistent stiffness-induced adaptations after exposure to 24-hours of increased culture surface stiffening, but not with 6 hours of stiffening.^11^ These stiffness-induced mechanical changes involve cytoskeleton-mediated mechanotransduction with persistence requiring induction of the post-translational detyrosination of a-tubulin^10^. While the role of cytoskeletal elements in mediating MM has been well characterized in our previous work^10^, the contribution of the nucleus in imprinting the stiffness-induced phenotype (MM), particularly in cardiomyocytes, has received limited attention to date. This gap in understanding presents an opportunity to explore how nuclear factors, such as the nuclear envelope and associated proteins might contribute to MM.

In the present study, we investigated the nuclear responses to transient increases in culture surface stiffness of durations sufficient to induce MM. To achieve this, we utilized a magnetorheological elastomer (MRE), which allows for controlled, bidirectional changes in stiffness, simulating conditions ranging from normal to diseased myocardium (9-56 kPa)^12,13^. We found that 6h of culture surface stiffening does not produce persistent cellular responses in isolated iPSC-derived cardiomyocytes. In contrast, 48h culture surface stiffening, sufficient to produce persistent increases of cell area, is associated with increased expression of lamin AC at the nuclear envelope, increased markers of DNA damage including double-strand breaks (DSBs), nuclear rupture events, and leaking of DNA repair factors to the cytoplasm. Inhibition of mechanotransduction to the nucleus by preventing post-translational a-tubulin detyrosination or inhibiting the LINC complex proteins connecting MTs to the nucleus, blocks MM, lamin induction, DNA damage and nuclear rupture. Conversely, prior knock-down of Lamin AC expression or inhibition of DNA repair responses accelerates the induction of MM and DNA damage markers. These studies link MM to DNA damage in cardiomyocytes for the first time, demonstrate an increased sensitivity to MM and DNA damage in the presence of lamin deficiency, and suggest that interventions to limit nuclear mechanotransduction or augment DNA damage responses might be able to prevent or delay the onset of MM during sustained increases in myocardial stiffness.

## Methods

### Magnetorheological elastomer (MRE) preparation

A previously described magnetorheological elastomer (MRE) model^12^ (Fig. 1A) was used to provide instantaneous and reversible stiffening and softening to study the responses of ventricular human iPSC-derived cardiomyocytes (NCardia, Inc.M068 Ncytes TM) and adult rat and human CMs to disease-relevant stiffness increments in the absence of cell-cell interactions. Briefly, the MRE is composed of a soft elastomer Sylgard 527 prepared per manufacturer’s directions by mixing equal weights of part A and part B then thoroughly mixed with Carbonyl Iron Powder (CIP-CC, BASF, Ludwigshafen, Germany) at a 1:1 mass ratio. 5 g of the mixture was then poured into 35 mm culture dishes, degassed for 10 min, and baked overnight at 60 °C. A magnetic field was applied to the culture dish through use of a cylindrical neodymium rare earth magnet (1–1/4″ ×.”, CMS Magnets, Garland, TX, USA).

**Figure 1:**
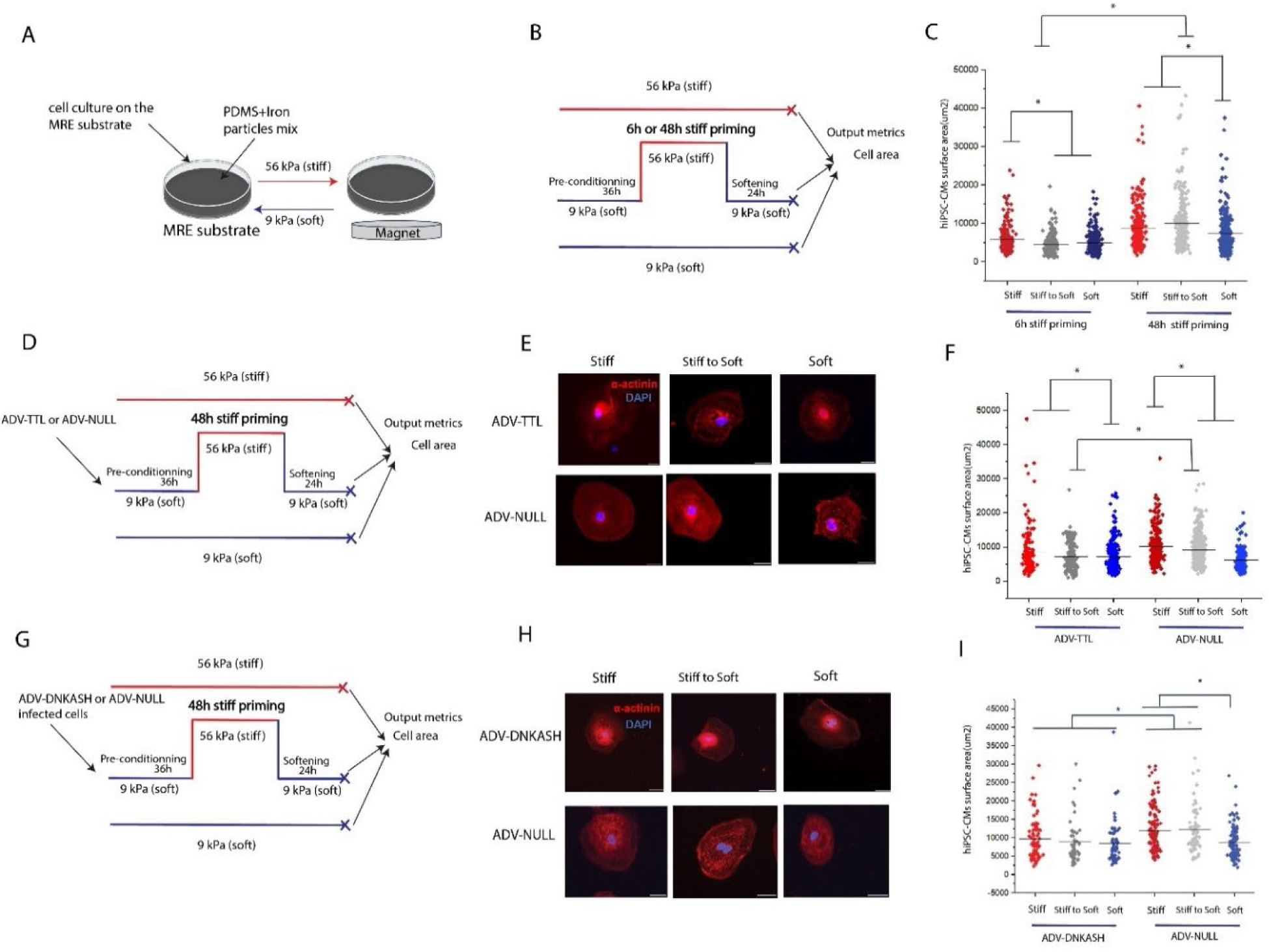
Mechanical memory induction in hiPSC-CMs is dependent on MTs and the LINC complex. A) Schematic of the cell culture substrate with tunable bidirectional capabilities, B) Schematic of the persistent stiffness induced phenotype experimental setup showing the 6h compared to the 48h priming stiff priming and after resoftening where MM is observed, C) hiPSC-CMs surface area in the 6h and 48h stiff priming conditions, following resoftening and in the stiff and soft control conditions D) MM experimental setup with the timepoint of ADV-DNKASH infection, E) Immunofluorescence images of actinin in hiPSC-CMs for the ADV-DNKASH and the respective control (ADV-NULL) F) ADV-DNKASH and ADV-NULL cell area comparison, G) MM experimental setup with the timepoint of ADV-TTL infection and H) Immunofluorescence images of a-actinin in hiPSC-CMs for the ADV-TTL and the respective control (ADV-NULL) I) Surface area for stiff, stiff to soft and soft for the ADV-TTL cells compared to their respective control (ADV-NULL).

### Induced pluripotent stem cell-derived cardiomyocytes (hiPS-CMs)

Human hiPS-CMs (NCardia, Inc.M068 Ncytes TM) were cultured following the protocol provided: Fibronectin was suspended in cold 4°C PBS 1:200 dilution mixed gently, and 5ml of this suspension was added to two T25 cell culture plates (Corning Catalog #) and incubated for 1hr at 37C to allow fibronectin to coat the surface. The solution was gently aspirated and 1 vial of human iPS cells were added to each plate in RPMI medium containing 10 μM of ROCK inhibitor (Y27632, #50-863-7-Fisher). Culture medium was replaced every other day (no ROCK inhibitor) until ready for the experiments.

### Immunofluroescence Staining

At the end of each experiment, hiPSC-CMs or adult rat and human CMs were rinsed with PBS and fixed in prechilled methanol at −20°C for 10 minutes. Cells were then washed three times with PBS and placed in blocking buffer (3% BSA in PBS) for 1 hour at room temperature. Cells were labeled with primary antibodies in 1% BSA overnight at 4°C either with anti-α-actinin (Sigma Aldrich #A7732), anti-phospho h2ax (ThermoFischer, #SAB5700329) or anti KU80 (Cell Signaling #2753S). Cells were then washed and labeled with secondary antibodies (555 Alexa Fluor goat anti-mouse) and (488 Alexa Fluor goat anti-rabbit) at 37°C for 1 hour. Finally, DAPI was added for 10 minutes in a dark environment. After a final wash with PBS, cultures on the 2D MRE were maintained in PBS until imaging.

### Cell area Quantification

Immunofluorescence images were collected on an upright microscope (Nikon eclipse 80i) with a 40x water-immersion objective. Each cell was individually measured for cell area using ImageJ. Regions of interest containing the entire cell excluding the nuclei were used to determine fractional coverage of cytoskeletal proteins. For hiPSC-CMs cell area was determined using α-actinin cytoskeleton. Area coverage was determined by a mask defining the cell border.

### DNA damage Quantification

Immunofluorescence images of the γ-h2ax collected on an upright microscope with a 40x water-immersion objective. Each cell was individually measured for γ-h2ax intensity using CellProfiler Software. The software uses object detection to detect all the nuclei with the γ-h2ax channel and quantifies the intensity of the DNA damage.

### Nuclear leakage of KU80 quantification

To quantify the extent of nuclear rupture and the leakage of DNA repair factors KU80 to the cytoplasm, the intensity ratio of cytoplasmic KU80 over nuclear KU80 levels were quantified using ImageJ by creating a mask to define the nucleus and the cytoplasm and measuring the intensity separately then calculating the ratio.

### Nuclear Morphometric Analysis

We used an analytic tool to quantify the number of cells in a population that present characteristics of senescence, apoptosis or nuclear irregularities through nuclear morphometric analysis. The tool previously described^14^ uses nuclear image analysis and evaluation of size and regularity of adhered cells in culture. From the different conditions, measurements of nuclear morphometry, principal component analysis filtered four measurements that best separated regular from irregular nuclei. These measurements, namely aspect, area box, radius ratio and roundness were combined into a single nuclear irregularity index (NII). Normal nuclei are used to set the parameters for a given cell type, and different nuclear phenotypes are separated in an area versus NII plot. Nuclear Morphometric Analysis (NMA) was determined based on a direct and objective way of screening normal, senescent, apoptotic and nuclear irregularities using image J and NMA plugin

### Pharmacological interventions

Pharmacological interventions were performed using 5 μM ivermectin (Sigma, I8898) for 24h to inhibit nuclear import of DNA repair factors,). At the end of drug treatment, cells were washed with PBS and fresh media was provided until the end of the experiment

### Cytoskeleton interventions

Intervention of the cytoskeletal components were performed using a genetic approach to prevent α-tubulin tyrosination by infecting cells with adenovirus encoding tubulin tyrosine ligase (TTL)^1015^

### RNA sequencing

To measure gene expression profiles in the different conditions, Two MREs for each condition were required to produce a sufficient quantity and quality (RIN > 6.0) of mRNA. The mRNA was extracted using the Zymo Quick RNA MicroPREP isolation kit (ZYMO RESEARCH, catalog R1050) following manufacturer’s protocol. Three conditions in duplicate were produced for each condition. For quantitative analysis, the Nanodrop spectrophotometer was used to measure the RNA concentration and quality. Quality was indicated by measuring the absorbance ratios (260/280). Ratios around 1.8 were classified as pure and therefore acceptable for gene sequencing. The mRNA was shipped to an external company (AZENTA) for next generation Sanger sequencing.

### RT-qPCR

At the end of the experiments, mRNA from CMs from each condition was isolated using Zymo Quick RNA MicropREP isolation kit and reverse transcribed to cDNA using qScript (Quantabio, catalog 95048). Quantitative PCR was then performed with SYBR Green reagents (Life Technologies, catalog S7563), with the primers given in Table 1 used to amplify targets. Tests were conducted in triplicate for each condition.

**Table 1.** summarizes the top 10 most differentially expressed genes in the stiff compared to the softcondition based on the adjusted p-value.

| Gene ID | Gene name | Expression |
| --- | --- | --- |
| ENSG00000019582 | **CD74** | Downregulated |
| ENSG00000162511 | LAPTM5 | Downregulated |
| ENSG00000140968 | IRF8 | Downregulated |
| ENSG00000255833 | **TIFAB** | Downregulated |
| ENSG00000020633 | **RUNX3** | Downregulated |
| ENSG00000100079 | LGALS2 | Downregulated |
| ENSG00000259498 | **TPM1-AS** | Upregulated |
| ENSG00000141506 | **PIK3R5** | Downregulated |
| ENSG00000133808 | MICALCL | Upregulated |
| ENSG00000162804 | SNED1 | Upregulated |
| ENSG00000154175 | ABI3BP | Upregulated |

### SiRNA knockdown of Lamin AC in hiPSC-CMs

hiPSC-CMs were plated in a 6-well plate at 200 k per well until they reached 80% confluency. A mixture of transfection reagent (Santa Cruz, sc-29528) and transfection media (Santa Cruz, sc-36868) was added to the transfection solution, and then to the cells following manufacturer’s protocol. LaminAC siRNA (Santa Cruz, sc-44805) was added to the transfection solution and next to the cells for 6 h at 37C. RPMI media was then added to the transfection media for 24h. Then, the transfection solution with RPMI mix was removed and replaced by fresh RPMI for 72 h prior to seeding. Transfection efficiency was verified by western blotting: cells were lysed in UREA buffer. Membranes were blocked and incubated with anti-Lamin AC (1:100, Santa Cruz, sc-166871), and then visualized with secondary antibodies (Donkey anti-Mouse, 926-68072, and Donkey anti-Rabbit, 926-32213, LI-COR). Blots were quantified by measuring band intensity using ImageJ and normalizing relative expression to the levels of GAPDH

### Protein lysate and Western Blot

After washing the cells with PBS, UREA buffer (8M UREA, Water, Glycerol, 1M DTT, 1.5M Tris, 20% SDS, PMSF) supplemented with 1x protease inhibitor cocktail (Cell Signaling Technology: 5872S) was added to the MRE substrate. Residual undissolved cell debris were removed from the resulting samples by centrifugation at 8000g for 10 minutes at room temperature (22°C). All samples were diluted to 4 µg/µL. The diluted total protein lysates were aliquoted and stored at −80°C until further processing. To quantify the relative abundance of specific proteins of interest in the total protein lysate, aliquoted diluted total protein lysate samples were thawed at room temperature. 4 µL of 4x loading buffer was mixed with 16 µL of the total protein lysate. The resulting samples were heated at 100°C for 10 minutes. The heated samples were vortexed thoroughly and loaded (15 µL/sample) onto precast protein gels (Bio-Rad: 5671085). Protein gel electrophoresis was carried out under constant current. The resolved proteins were transferred onto a nitrocellulose membrane using the Turbo Transfer System (Bio-Rad) under recommended conditions. The post-transferred membrane was blocked in blocking buffer (3% in PBST) for at least 1 hour at room temperature. The blocked membrane was incubated overnight at 4°C with primary antibodies (anti γ-h2ax ThermoFischer, SAB5700329, anti-KU80, Cell Signaling #2753S and anti-Lamin AC, Santa Cruz, sc-166871) diluted in 1x Tris buffered saline. The membrane was washed twice using TBST and incubated for 1 hour at room temperature with secondary antibodies (Donkey anti-Mouse, 926-68072, and Donkey anti-Rabbit, 926-32213, LI-COR). The final immunoblotted membrane was washed twice using TBST and was imaged using the Odyssey Western Blot Imaging System (LI-COR Biosciences).

### Experimental replicates and statistical analyses

For all experiments, a minimum of three biological replicates were performed for each condition and a minimum of 100 cells were analyzed for each replicate. All data are presented as the mean±SEM. For most experiments, comparisons were made between data from culture surfaces that were persistent stiff, persistently soft, and transiently (stiff-to-soft) with varying durations of stiffening before re-softening (see Fig 1). For the experimental metrics defined above, two-way ANOVA was used for studies comparing multiple time points across these three experimental conditions. For all statistical tests, a *p*-value of <0.05 was considered significant.

## Results

### Mechanical memory induction in hiPSC-CMs is dependent on MTs and LINC complex

The primary advantage of our MRE model is its ability to precisely and reversibly control culture surface stiffness. (Fig. 1A). Our initial experiments were guided by our prior studies identifying the duration of culture on the stiffened surface (stiff priming with magnet) required for phenotypic adaptations to persist after restoration of the softened culture conditions (magnet removed). After hiPSC-CMs underwent an interval of 36 hours of soft preconditioning on a culture surface of 9 kPa stiffness, the culture surface was stiffened for either 6h or 48h prior to a restoration to the 9 kPa stiffness (Fig 1B). In these experiments, cell profile area was the primary indicator of mechanoresponses using immunofuorescence staining of a-actinin. We found that stiff priming for 6h followed by resoftening led to a return to baseline (soft) cell profile area. However, stiffening for 48h led to a persistent increase in cell area even after resoftening indicating induction of MM (Fig.1C)

Next, we wanted to investigate the role of the cytoskeletal components with a specific focus on the MT network on the induction of MM. We used ADV-TTL to suppress the detyrosinated MTs prior to stiff priming for 48h (where MM is induced) (Fig 1D). As a control, ADV-NULL cells were seeded on the MRE substrate for the same duration. We found that suppressing the detyrosinated MTs did not prevent the increase of iPSC-CMs area during 48 h of stiff priming, however cell area returned to the soft condition levels upon resoftening (Fig 1E-F). This indicates that MM induction in hiPSC-CMs is dependent on detyrosination of a-tubuliin.

To further probe the mechanotransduction pathway required to imprint MM, we disrupted the linker of nuclear skeleton and cytoskeleton (LINC) complex. We transfected hiPSC-CMs with a dominent negative KASH domain construct which is known to effectively uncouple the cytoskeleton from the nucleus^15^. Infected cells were cultured on the soft MRE for 36 hours before the cells were stiff primed for 48h (Fig 1G). ADV-NULL (control) cells were also seeded on the MRE substrate for the same duration. Persistent soft and persistent stiff culture substrates served as controls for both the ADV-DNKASH and ADV-NULL cells. We observed that increases in cell area observed with 72 hours of culture substrate stiffening or stiff priming for 48 hours followed by resoftening (Fig 1H-I) in ADV-NULL infected cells are completely blocked in cells with LINC complex disruption via ADV-DNKASH. These data indicate that mechanical forces applied to the cell surface are transmitted through the cytoskeleton and are then conveyed to the nucleus via the LINC complex.

### Lamin AC expression adapts to conditions associated with MM

Recognizing the ability of the nucleus to sense and respond to mechanical forces and recognizing the role of the nuclear lamina as part of the mechanotransduction pathway, we wanted to explore how lamin AC levels will respond to the increase in stiffness after both 6h and 48h of stiff priming and following 24h of resoftening. All experiments were performed similar to the previous protocols (Fig 2A) and Lamin AC levels were quantified using western blot at the end of the stiff and soft cultures for both 6h and 48h priming, and at the end of the resoftening phase. As shown in Fig. (2B-D), we observed that 6h of stiff priming was insufficient to induce an increase in the levels of Lamin AC expression across all conditions. However, stiff priming of 48h consistently induced an increase of Lamin AC levels compared with the control culture on soft surface (9 kPa). When the magnet was removed after 48 hours to allow a return to 9 kPa, the levels of Lamin AC remained high (Fig 2B and E-F). Extending these findings, we measured levels of Lamin AC in iPSC-cardiomyocytes with LINC complex disruption via ADV-DNKASH (Fig 2G). After LINC complex disruption, we found that Lamin AC levels did not increase in response to the increase in stiffness (Fig 2H-I). These results demonstrate that the nuclear envelope responses to mechanically transduced extracellular stiffening signals include upregulation of Lamin A/C expression, and that this upregulation requires an intact LINC complex and more than 6 hours to be evident at the protein level.

**Figure 2:**
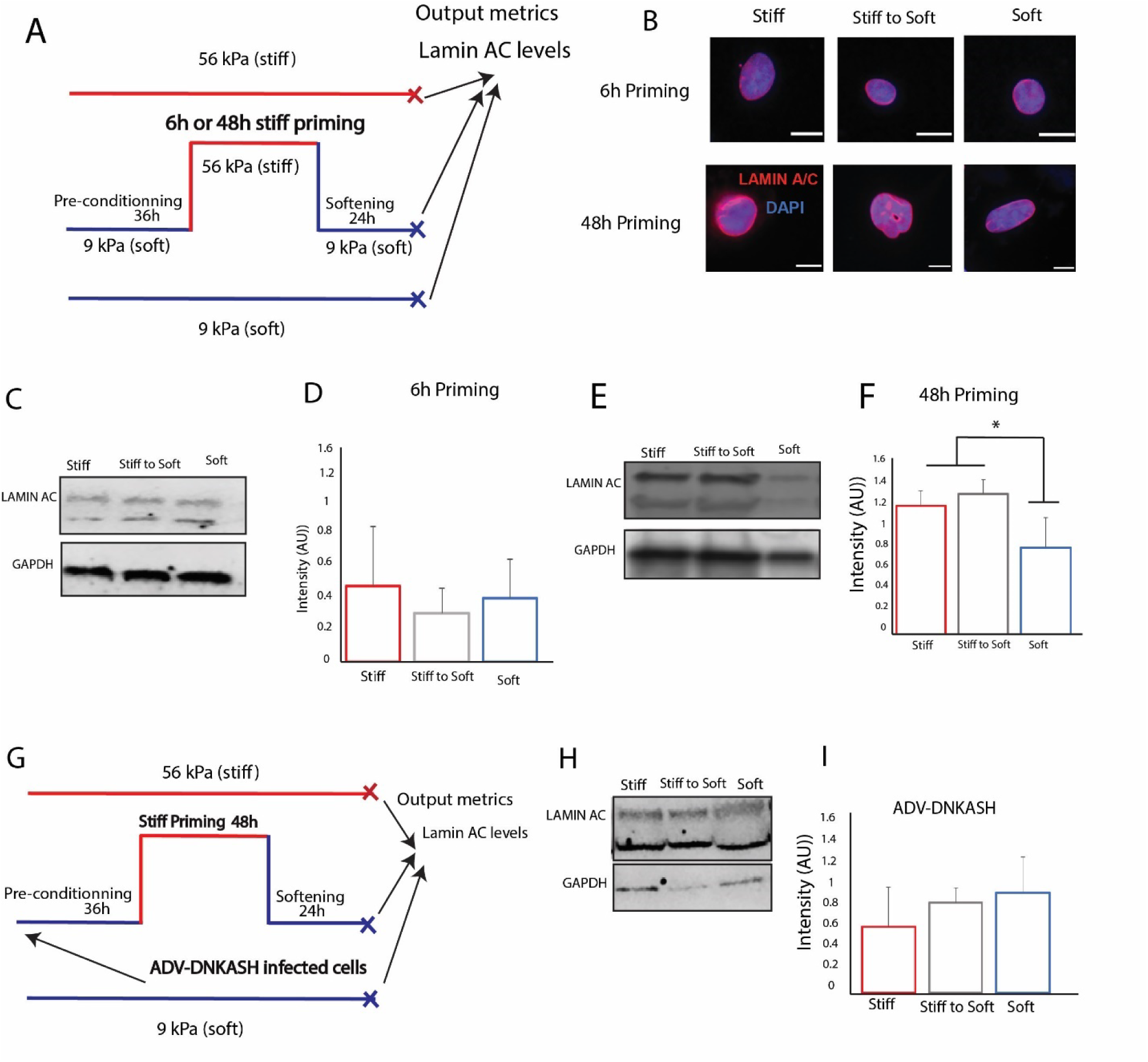
Lamin AC expression adapts to conditions associated with MM. A) MM experimental setup with with Lamin AC levels as an output metric B) Immunofluorescence images of the nucleus in iPSC-CM showing Lamin AC localization for stiff, stiff to soft and soft for the 6h and 48h stiff priming respectively. C)Western blot of Lamin AC expression in the 6h priming D) Expression levels of Lamin AC using western blot under 6h of stiff priming, E) Western blot of Lamin AC expression in the 48h priming, F) Expression levels of Lamin AC using western blot under 48h of stiff priming, G) MM experimental setup with the timepoint of ADV-DNKASH infection with Lamin AC levels as an output metric, H) Western blot of Lamin AC expression and,and E) quantification of Lamin AC expression in the ADV-DNKASH cells under stiff, stiff to soft and soft conditions. Scale bar, 2 µm. Bar graphs depict mean±SEM and * depicts p<0.05.

To further understand nuclear mechanotransduction, we performed Bulk RNA sequencing on hiPS-CMs for the stiff priming associated with MM (48h of stiff priming), cells from three replicates of this experiment were pooled and then subjected to RNA-seq analysis and compared to persistent soft (Supplementary Fig 1A). Compared with culture on the persistently soft surface, there were 25 significantly upregulated genes and 4 downregulated genes after 72h on the stiffened culture surface. Among the 29 differentially expressed genes in the stiff condition compared to the soft, there were approximately 40% genes that have been associated with DNA damage or DNA repair (Table 1). Specifically, genes linked to DNA damage (e.g. TPM1)^16^ are upregulated (Supplementary Fig B) and genes associated with DNA repair (e.g. RUNX and CD74) ^17^ are downregulated in cells cultured for 48 hours on the stiffened surface. These findings suggest that pathological extracellular stiffness conditions induce a cell-autonomous downregulation of the DNA repair mechanisms in hiPSC-CMs that are otherwise active under soft culture conditions, potentially contributing to the activation of genes associated with DNA damage.

We next focused on how the duration of stiff priming affects DNA damage and repair factors using RT-qPCR and the reversibility of gene expression changes induced by transient stiffening. All experiments began with preconditioning the cells on soft MRE for 36h, followed by an increase in culture surface stiffness for either 6 or 48 hours followed by a return to the softened culture surface (9 kPa) for another 24 hours via magnet removal. After the MRE had been restored to a 9 kPa stiffness for 24 hours mRNA was extracted from the hiPSC-CMs. As controls, hiPSC-CMs were either continuously cultured on stiff MREs (56 kPa) or on soft MREs (9 kPa) for the entire duration of the experiments beginning at the end of the preconditioning phase. After 48 hours of stiff priming and 24 hours of softening, RT-qPCR analysis demonstrated that RUNX3 and CD74 were downregulated compared to the control soft condition (Supplementary Fig 1D). However, 6 hours of stiff priming followed by 24 hours of resoftening led to an *upregulation* of RUNX3 expression, suggesting the potential involvement of DNA repair factors in mitigating DNA damage following short durations of stiff priming, with a failure to maintain this response with 48 hours of stiff priming.

### MM conditions are associated with DNA damage and nuclear rupture

To examine the functional impact of changes in genes associated with repair of DNA damage, we quantified levels of phospho-histone γ-h2ax, an established marker for detecting DNA damage^18^ in the 6h and 48h stiff priming conditions, and following resoftening 24h of resoftening (Fig. 3A). As shown in Fig (3B-D) and Supplementary Fig 2 A, after 6 hours of stiff priming, we did not observe any significant change in the levels of DNA damage between the stiff, stiff-to-soft, and soft conditions. However, 48 hours of stiff priming induces an increase in DNA damage foci in the nucleus which remain evident after 24 hours of resoftening (Fig 3B and 3E-F and supplementary Fig 2B). In addition, we found that 48h of stiff priming also induces an increase in the size and number of DNA damage foci (supplementary Fig 2 C-D) which was not significant in the 6h priming duration. A closer inspection of DAPI stained nuclei using the NMA tool revealed nuclear abnormalities in the 48h priming condition on the stiff substrate and following resoftening with increased irreversible and abnormal nuclear area (Supplemental Fig 3 and 4). Resoftening after 48h of stiff priming didn’t recover nuclear abnormalities with more than 19% cells appearing irregular, senescent and apoptotic, whereas approximately 98% of the cells in the 6h priming condition appeared normal in size and shape. This increase in apoptotic and senescent cells in the 48h priming condition was present despite activation of the DDR pathway.

**Figure 3:**
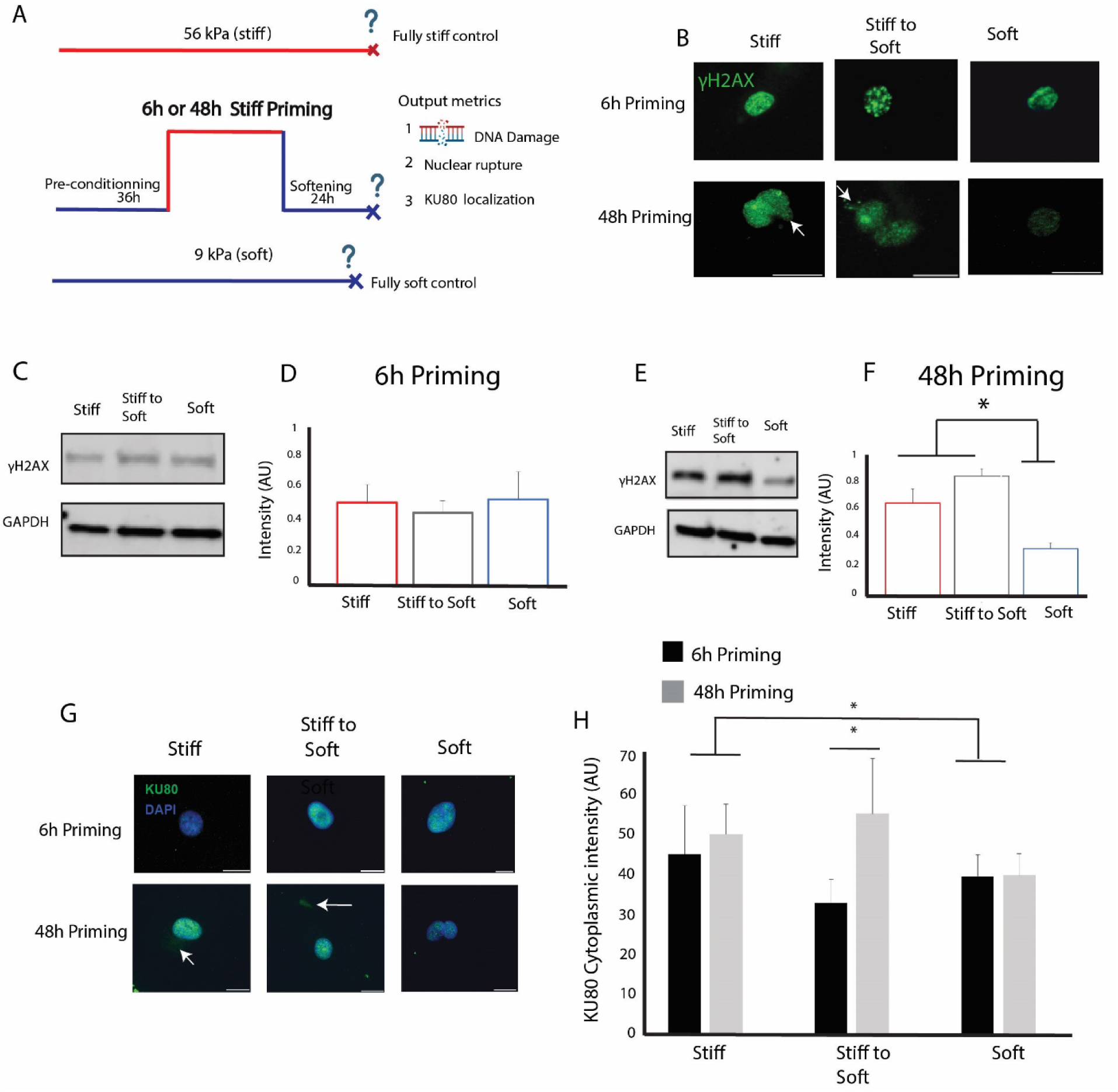
MM conditions are associated with DNA damage and nuclear rupture. A) Schematic of the persistent stiffness induced phenotype experimental setup showing the 6h compared to the 48h priming and after resoftening, with the time points of DNA damage measurement, B) Immunofluorescence images of the nucleus in iPSC-CM showing y-h2ax localization for stiff, stiff to soft and soft for the 6h and 48h stiff priming respectively, C)) Expression levels of γ-h2ax using western blot for 6h of stiff priming, D) Verification of the γ-h2ax levels using western blot under 6h of stiff priming, E) Expression levels of γ-h2ax using western blot under 48h of stiff priming, F) Verification of the γ-h2axlevels using western blot under 48h of stiff priming, G) Immunofluorescence images of the KU80 localization in the cells for the 6h and 48h stiff priming conditions and, H) Quantification of the mislocalization of the KU80 to the cytoplasm in the 6h and 48h priming conditions. Scale bar, 5 µm. Bar graphs depict mean±SEM and * depicts p<0.05.

**Figure 4:**
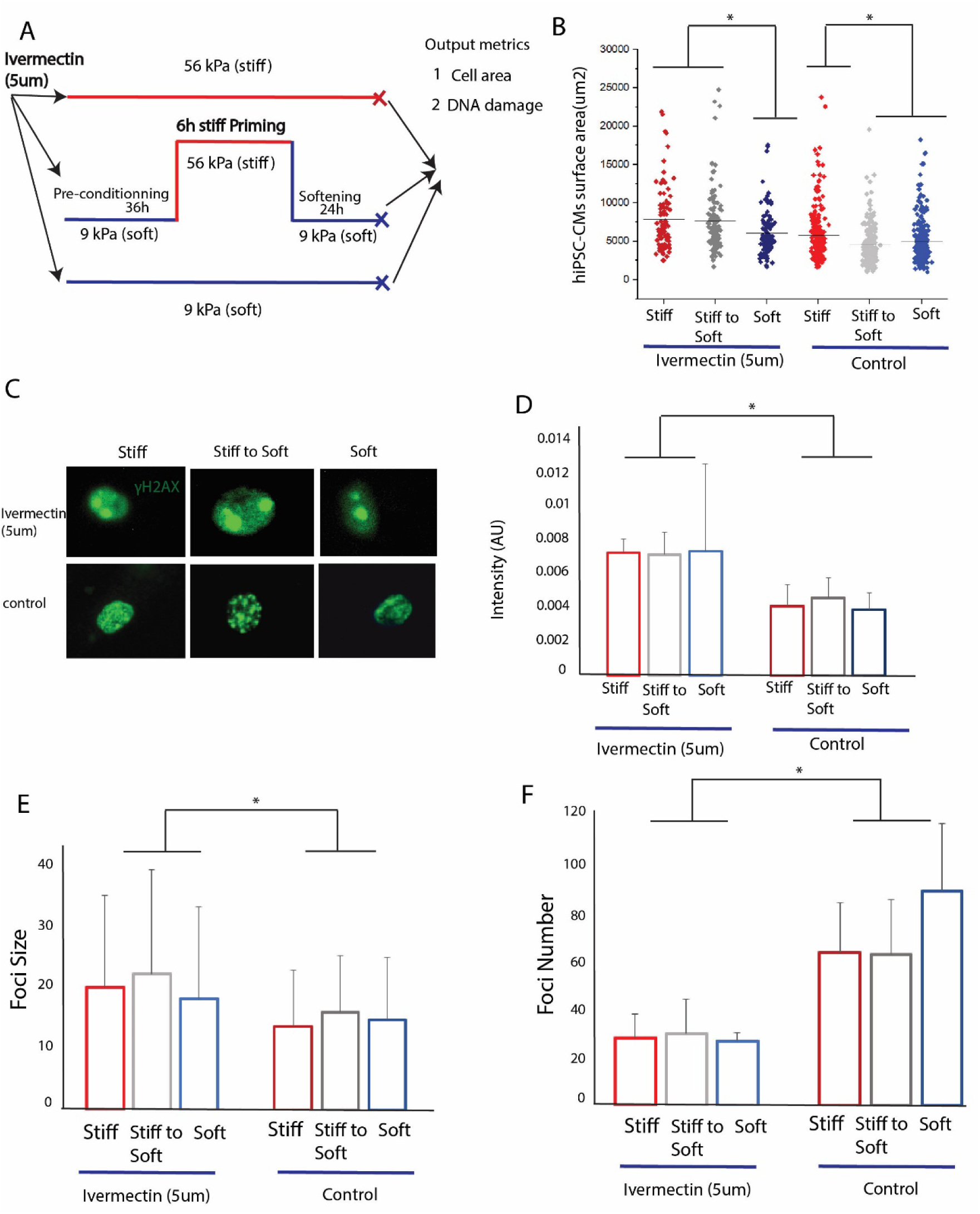
The role of the DDR mechanism in imprinting or reversing mechanical memory. A) Schematic of the cell culture experiment where Ivermectin was added B) area for stiff, stiff to soft and soft for the ivermectin treated cells, C) Immunofluorescence images of γ-h2ax in control and Ivermectin treated cells following 6h of stiff priming and following resoftening and soft control, D)) Quantification of the γ-h2ax levels using IF intensity in the 6h priming condition following ivermectin treatment compared to the control, E) Quantification of DNA Damage Foci size and F) number in the 6h priming condition with ivermectin and control for stiff, stiff to soft and soft conditions,

Based on immunofluorescence images (Fig. 3B), we also found that γ-h2ax foci were leaking outside the nuclear membrane after 48h of stiff priming, but not after 6h of stiff priming. This indicates that mechanical stress durations associated with MM not only induce DNA damage but may also compromise nuclear integrity. Accordingly, we next explored the extent of nuclear rupture by measuring the presence of DNA repair factors in the cytoplasm using the cytoplasmic mis-localization of KU80 as a readout. We found that the DNA repair protein KU80 was mis-localized to the cytoplasm after 48h of stiff priming and following resoftening compared to the 6h priming (Fig 3G-H), suggesting that escape of DNA repair factors from the nucleus might be linked to irreversibility of cellular responses to extracellular stiffening whereas nuclear retention of DNA repair factors might favor reversibility of cellular responses.

### Reversing the persistent stiffness induced phenotype in CMs is dependent on the DNA repair factors and the activation of DNA damage response (DDR)

To further explore how localization of soluble DNA repair factors might affect MM induction, we treated cells during the pre-conditioning phase with 5uM of Ivermectin a known nuclear transport inhibitor that blocks nuclear import of KU80 in cytoplasm^19^. In experiments utilizing only 6h of stiff priming prior to resoftening of 24h (Fig 4A), we found that inhibiting the nuclear import of DNA repair factors speeds up MM induction in hiPSC-CMs, evidenced by the irreversible change in cells area after resoftening compared to untreated control cells, as shown in Fig 4B. We also found that blocking the import of the DNA repair factors increased DNA damage levels compared to the control conditions (Fig 4C-E). In addition, we noticed that ivermectin increased the size of DNA damage foci (Fig 4F) in all conditions (stiff, stiff to soft and soft). Nuclear morphology analysis also revealed an increase in irregular nuclei after resoftening compared to the respective control (Supplemental Fig 6) for the 6h priming duration with ivermectin consistent with an increase in the percentage of senescent cells. Together, these results indicate that proper nuclear import of DNA repair factors within stressed nuclei is required to prevent DNA damage and distortions of nuclear architecture, under culture conditions when MM would not otherwise occur.

### Reduced Lamin AC expression accelerates MM and DNA damage during short duration stiffness challenges

To explore the functional impact of increased lamin AC expression on limiting DNA damage and the link between DNA damage and imprinting MM, we employed siRNA mediated knockdown Lamin AC levels in hiPSC-CMS prior to stiff priming. (Fig 5A). Depletion of LaminA/C was confirmed by western blot, with an approximately 50% reduction in Lamin A/C at 4 days post transfection compared to control cells (Fig 5B-C). We assessed the effects of lamin A/C reduction in iPSC-CMs together with a 6-hour interval of stiff priming that would not ordinarily be associated with DNA damage or MM (Fig 5D) Based on cell surface area changes, MM was induced in Lamin AC depleted cells, but not control cells with scrambled siRNA, after 6h of stiff priming and cells area increased and did not reverse after resoftening (Fig 5E). In addition, lamin A/C depletion substantially increased the degree of DNA damage (based on γ-h2ax) observed with persistent culture surface stiffening for 30h, with a more variable effect on the DNA damage markers after 6h stiff priming followed by 24h resoftening (Fig. 5F-G). These results indicate that upregulation of Lamin AC during extracellular stiffening likely contributes to the more sustained time required to elicit MM in iPSC-CMs and serves to limit the time-dependent DNA damage associated with exposure to extracellular stiffening.

**Figure 5:**
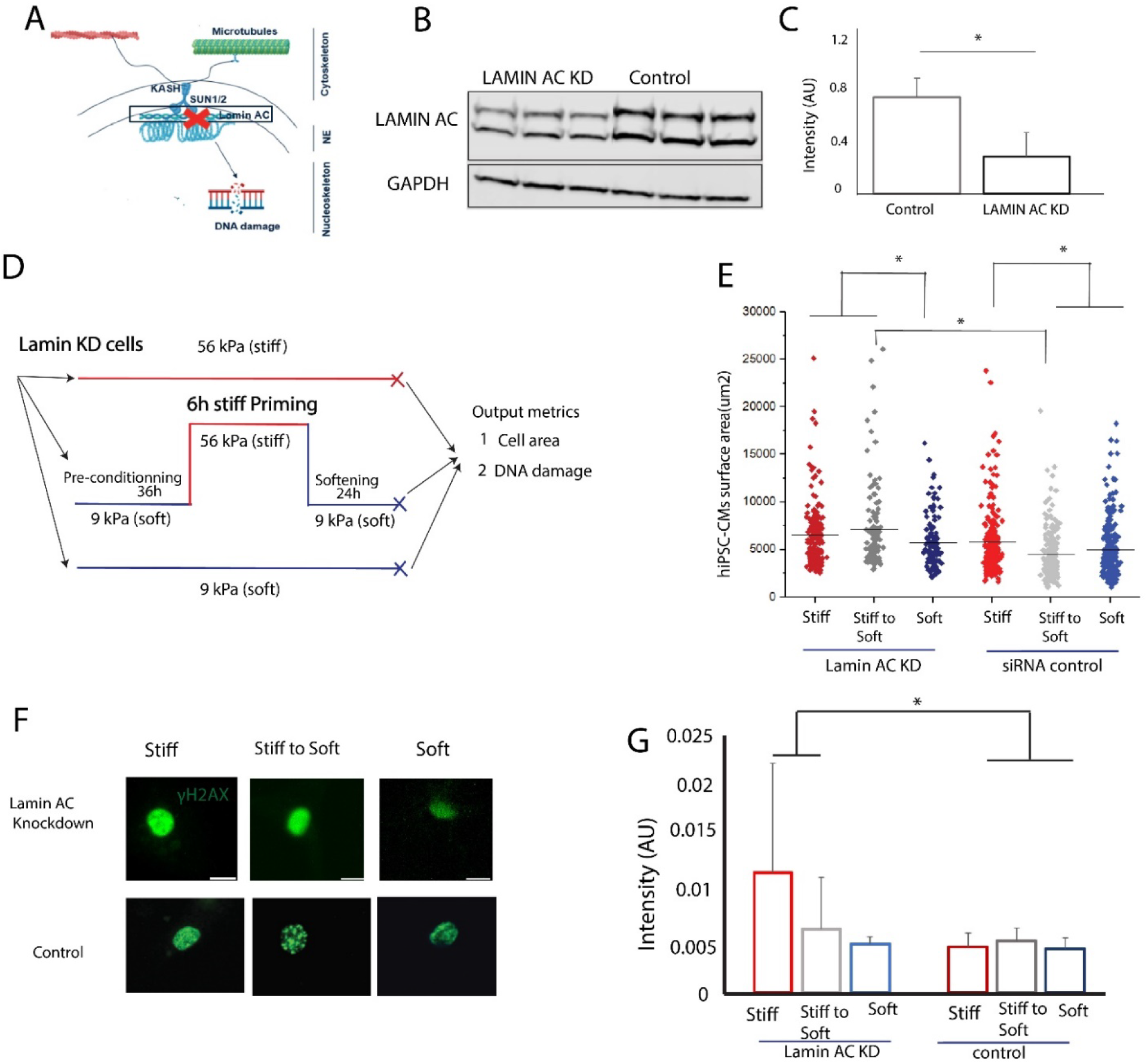
Lamin AC expression lessens MM and DNA damage during short duration stiffness challenges. A) Molecular conduits of mechanical memory with Lamin AC location B) Verification of the siRNA knockdown of Lamin AC using western blot under 48h of stiff priming, C) Quantification of Lamin AC levels in the control and knockdown, D) Experimental setup for MM induction in Lamin AC knockdown cells, E) hiPSC-CMs surface area for the stiff, stiff to soft and soft respectively following 6h of stiff priming and 24h of resoftening, F) Immunofluorescence images of the γ-h2ax marker in Lamin AC KD cells and control, G) Quantification of DNA damage in Lamin AC KD cells and scrambled control cells. Scale bar, 2 µm. Bar graphs depict mean±SEM and * depicts p<0.05.

### Stable MT network increases nuclear tension to promote DNA damage and persistent stiffness induced phenotype

In prior studies, we demonstrated that microtubules (MT) play an essential role in transducing iPSC-CM responses to extracellular substrate stiffening, and that post-translational a-tubulin detyrosination is necessary for induction and maintenance of MM in iPSC-CMs^10^ (Fig 1). To investigate whether MT detyrosination is required for stiffness-induced DNA damage, we infected cells with ADV-TTL to suppress detyrosination prior to 48h of stiff priming followed by resoftening for 24 hours (Fig. 6A). We found that ADV-TTL-infected cells had lower levels of DNA damage based on γ-h2ax staining compared with ADV-NULL-infected cells (Fig. 6B-C). ADV-TTL infection largely suppressed the formation of excess foci (Supplemental Fig 7A-B). These results suggest that suppression of detyrosinated microtubules during the preconditioning phase can reduce DNA damage associated with MM. To assess the relative importance of limiting MT-mediated force transduction vs. attenuating the DNA damage response, we suppressed detyrosinated microtubules with ADV-TTL in hiPSC-CMs in combination with Ivermectin treatment in 48h stiff priming experiments (Fig 6D). We observed that, even in the presence of ivermectin to inhibit the transport of KU80 back to the nucleus, ADV-TTL pretreatment allowed cell area to return to levels observed with a persistently soft culture surface (Fig. 6E and supplemental Fig 6C) and DNA damage is prevented (Fig 6F-G). This indicates that blocking nuclear mechanotransduction by suppressing a-tubulin detyrosination limits MM and DNA damage, even when the nuclear import of DNA repair factors is inhibited.

**Figure 6:**
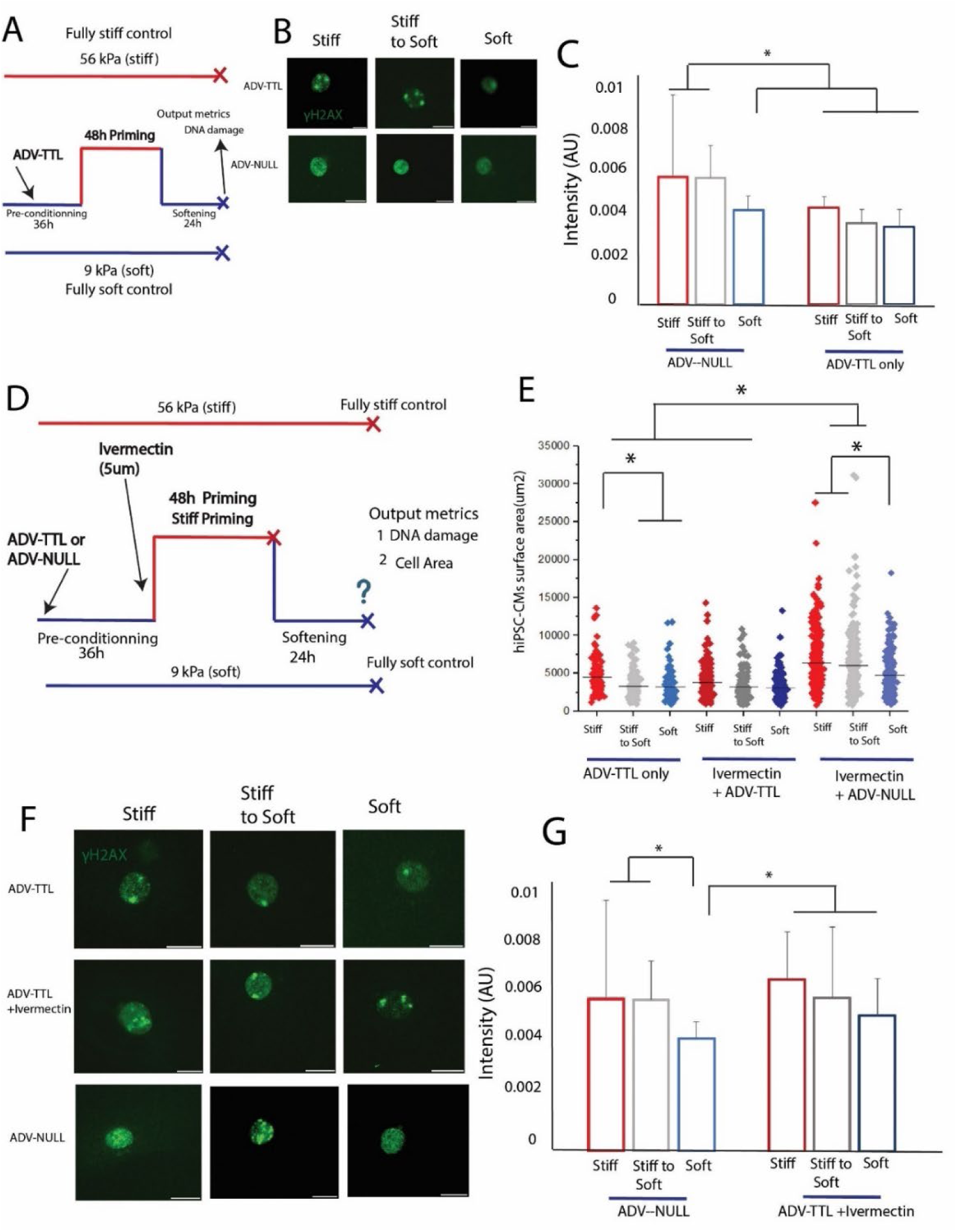
Stable MT network increases nuclear tension to promote DNA damage and persistent stiffness induced phenotype. A) Schematic of mechanical memory protocol with ADV-TTL infection before the 48h stiff priming phase, B) Immunofluorescence images of DNA damage in ADV-NULL and in ADV-TTL cells, C) Quantification of DNA damage intensity in ADV-NULL (MM) and) ADV-TTL cells, D) Schematic of ADV-TTL infection in the 48h priming condition with the ivermectin treatment with cell surface area and DNA damage as the output metrics, E), E) hiPSC-CMs surface area comparison between the ADV-NULL, ADV-TTL and the ivermectin+ ADV-TTL, F) Immunofluorescence images of γ-h2ax in hiPSC-CMs in the control (ADV-NULL), the ivermectin+ ADV-TTL and ADV-TTL only, G) Quantification of DNA damage levels in the ivermectin treated cells with ADV-TTL infection compared to only ADV-TTL cells. Green y-h2ax, Red a-actinin. Scale bar for DNA damage 2 µm. Bar graphs depict mean±SEM and * depicts p<0.05.

### The LINC complex is essential for force transduction to the nucleus and the initiation of DNA damage

To investigate the role of the nuclear envelope and the LINC complex in DNA damage, we used adenoviral infection with ADV-DNKASH to disrupt the physical connection between the cytoskeleton and the nucleus during the preconditioning phase. Following infection, cells were subjected to a stiff-priming protocol for 48 hours (Fig. 7A), followed by a 24-hour period of soft substrate exposure (resoftening). Compared to control cells infected with ADV-NULL, ADV-DNKASH-infected cells exhibited significantly reduced levels of DNA damage (Fig. 7B–C). Disruption of the LINC complex and subsequent nuclear decoupling effectively inhibited the formation of excess DNA damage foci. These findings support a critical role for the LINC complex in nuclear mechanotransduction and its contribution to mechanosensitive DNA damage responses

**Figure 7:**
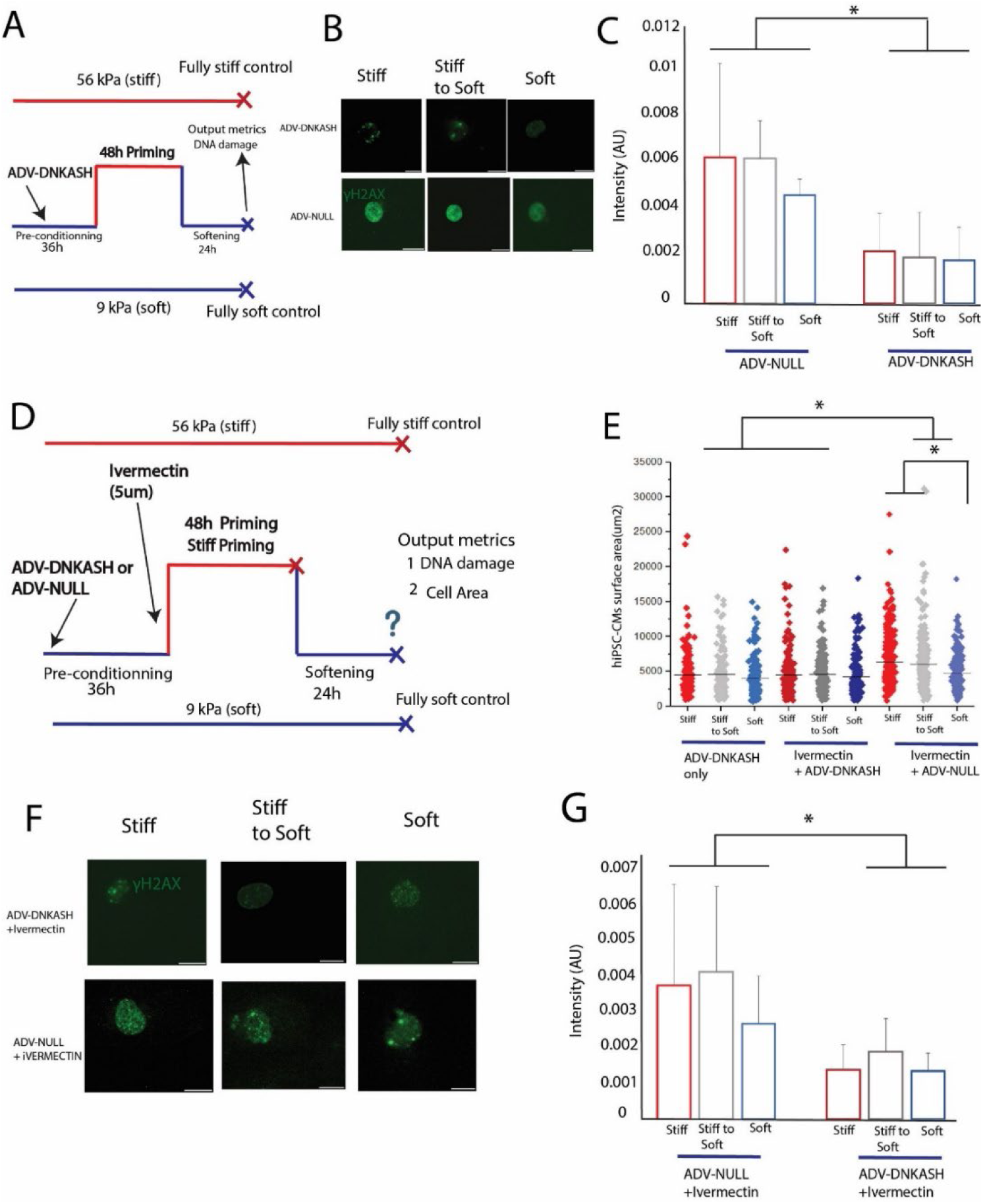
The LINC complex protects the nuclear tension to promote DNA damage and persistent stiffness induced phenotype. A) Schematic of mechanical memory protocol with ADV-DNKASH infection before the 48h stiff priming phase B) Immunofluorescence images of DNA damage in ADV-DNKASH and ADV-NULL cells, C) Quantification of DNA damage intensity in control (MM) and) ADV-DNKASH cells, D) Schematic of ADV-DNKASH infection in the 48h priming condition with the ivermectin treatment with cell surface area and DNA damage as the output metrics,, E) hiPSC-CMs surface area comparison between the control (ADV-NULL),the ivermectin+ ADV-DNKASH,and the ADV-NULL+ivermectin, F) Immunofluorescence images of γ-h2ax in hiPSC-CMs in theADV-NULL+ivermectin, and the ivermectin+ ADV-DNKASH, G) Quantification of DNA damage levels in the NULL control cells +ivermectin cells and, ADV-DNKASH+ivermectin. Green y-h2ax, Scale bar for DNA damage 2 µm. Bar graphs depict mean±SEM and * depicts p<0.05.

With ADV-DNKASH, cell area remained low despite cell surface stiffening indicating an impaired mechanosensitive response (Fig 7E).. Even in the presence of ivermectin (Fig 7D), ADV-DNKASH-infected hiPSC-CMs failed to respond to increased substrate stiffness.(Fig 7E-G and supplemental Fig 7).These results suggest that decoupling the nucleus from the cytoskeleton limits the transmission of mechanical stress that would otherwise lead to DNA damage. By attenuating nuclear stress, disruption of the LINC complex provides a protective mechanism against mechanically induced genomic instability.

## Discussion

The present study offers new insights into how nuclear mechanosensing and DNA damage and contribute to MM, defined as the persistence of phenotypic changes in hiPSC-CMs following a transient exposure to a biomechanical stimulus. Using a cell culture substrate with bidirectionally tunable stiffness, we investigated MM in hiPSC-CMs with a particular focus on nuclear mechanoresponses. Our findings demonstrate that culture on stiff substrates for 48 hours, followed by a restoration of baseline stiffness, is sufficient to induce persistent increases in cell size consistent with MM. These morphologic changes indicating MM are transmitted mechanically through MTs and the LINC complex. Our results indicate that extended mechanical stress and increased nuclear tension lead to increased nuclear irregularity, and numerous molecular signals marking induction of DNA damage and DNA repair responses with disruptions to the nuclear membrane despite upregulation of Lamin A/C expression. Disruption of the nuclear envelope allows leakage of DNA repair factors into the cytoplasm which exacerbates DNA damage. We observe that preventing the increase in Lamin A/C expression or constraining the nuclear import of DNA repair factors accelerates the induction of MM and DNA damage to as little as 6 hours. This suggests that these factors are crucial for preserving nuclear integrity under short-term mechanical stress. These findings demonstrate that increases in extracellular stiffness, transduced to the nuclear membrane by MTs, induces MM by inducing DNA damage responses. Our results are consistent with previous studies linking Lamin A/C dysfunction to DNA damage.^20,21^

These studies establish a consistent link between conditions that induce MM and molecular markers of DNA damage. While persistent DNA damage and activation of the DNA damage response (DDR) have been reported under pathological stressors such as oxidative and metabolic stress^22^ [PMID: 33911272], the role of DDR activation in cardiomyocytes has only been explored in a limited number of studies^23–27^. DDR signaling has been shown to contribute to pathological cardiomyocyte hypertrophy, with affected cells exhibiting impaired contractile and electrophysiological function, leading to cardiomyopathies and arrhythmias^20,28–30^. However, a direct correlation between DDR activation and mechanical memory in cardiomyocytes has not been reported previously.

DNA damage and cellular senescence are heightened in various disease-linked mutations, particularly those affecting the nucleoskeletal protein lamin-A (LMNA)^31,32^, and mutations of DNA repair factors, such as KU80^33^. Our studies underscore the crucial role of nuclear integrity and DNA repair factors in maintaining cellular function and reversing the stiffness-induced phenotype during short duration biomechanical stress. These findings also reveal the therapeutic potential of inhibiting DNA damage and activating the DDR pathway as disruptors of MM.

These studies also highlight the central role of MTs in mediating nuclear mechanotransduction and MM in cardiomyocytes. In our study, we found that suppressing a-tubulin detyrosination using ADV-TTL prior to culture on stiff MREs reduced DNA damage levels and permitted reversal of the stiffness-induced phenotype upon resoftening. Similarly, disrupting the interaction of MTs with the nuclear membrane by targeting the LINC complex with ADV-DNKASH also prevented MM and DNA damage. These findings indicate that mitigating the transduction of extracellular mechanical stimuli to the nucleus likely contributes to the reduction DNA damage. At the same time, beyond their well-known role in force transmission^34–37^, MTs are also essential for efficient DNA repair^34,38,39^, and our studies suggest that tyrosinated MTs, in particular, may favor a reparative DNA damage response (DDR). In this context, it is noteworthy that limiting a-tubulin detyrosination with ADV-TTL prevented MM and DNA damage even when nuclear import of DNA repair factors was blocked with ivermectin.

The heightened susceptibility to MM of iPSC-CMs with Lamin AC knockdown may have pathophysiologic significance. We observed that only 6 hours exposure to increased culture surface stiffness was sufficient to induce MM and DNA damage in the presence of reduced Lamin AC expression and a constrained ability to upregulate Lamin AC expression in response to the mechanical challenge. Such findings are consistent with the characterization of lamins as “shock absorbers” buffering the intranuclear propagation of mechanical stimuli reaching the nuclear membrane^40^.From this perspective, protecting the nucleus from increased mechanical stress by limiting a-tubulin detyrosination or disrupting the LINC complex are potential therapeutic strategies for cardiomyopathies associated with pathologic Lamin A/C variants.

### Limitations

While 2D cultures are well-suited for investigating cell-autonomous responses biomechanical responses, they lack the complexity of the in vivo environment where cells interact with one another and with the extracellular matrix in three-dimensional space. The complexity of tissue architecture in vivo likely alters how cells sense and respond to mechanical cues, especially when considering different cell types and their roles in the overall tissue response. These in vivo interactions might also affect the timing and mechanisms underlying DDR involvement in MM. Our use of hiPSC-derived human cardiomyocytes in these studies permitted sustained culture conditions required for adenovirus-medicated gene manipulation and 4-day MM protocols. However, these cells are relatively immature, and findings might differ adult cardiomyocytes. Our studies focused on the responses of the transition from a culture surface matching the stiffness reported for normal myocardium (9 kPa) and that reported for diseased myocardium (56 kPa)^12^. It is possible that the MM phenomena or timing observed might differ with different magnitudes of transient culture surface stiffening. Finally, most of our protocols allowed 24 hours of culture surface resoftening to define whether stiffness-induced changes were persistent, but it is possible that longer intervals of resoftening might permit some degree of reversibility despite the DNA damage and nuclear markers of senescence observed.

## Conclusions

In summary, these studies highlight the temporal aspects of stiffness-induced cardiomyocyte remodeling that are relevant to myocardial pathology. Short durations of extracellular matrix stiffening trigger reversible cardiomyocyte adaptations, while longer duration exposures induce MM that is consistently associated with markers of DNA damage, despite increases in nuclear membrane lamin expression. Acceleration of MM induction by lamin knockdown suggests that hereditary laminopathies may be associated with increased cardiomyocyte vulnerability to mechanical stress inducing DNA damage and senescence. Conversely, the protective effects of limiting stiffness-induced a-tubulin detyrosination, or attenuating mechano-transduction to the nucleus via the LINC complex, suggest potential cardioprotective strategies in the setting of laminopathies and/or sustained increases in extracellular matrix stiffness.

## Contributions

N.B. and K.B.M. developed the research strategy, N.B. and K.B.M. participated in the design of the experiments. N.B, S.P. and V.S. performed experiments. N.B. and K.B.M. participated in the writing of the manuscript. All authors participated in the review of the manuscript.

## Acknowledgement

We would like to extend our sincere gratitude to Professor Benjamin Prosser for generously providing us with the ADV-TTL and ADV-DNKASH viruses.

